# Electric Egg-Laying: Effect of Electric Field in a Microchannel on *C. elegans* Egg-Laying Behavior

**DOI:** 10.1101/2020.09.01.276097

**Authors:** Khaled Youssef, Daphne Archonta, Terrance J. Kubiseski, Anurag Tandon, Pouya Rezai

## Abstract

In this paper, the novel effect of electric field (EF) on adult *C. elegans* egg-laying in a microchannel is discovered and correlated with neural and muscular activities. The quantitative effects of worm aging and EF strength, direction, and exposure duration on egg-laying is studied phenotypically using egg-count, body length, head movement, and transient neuronal activity readouts. Electric egg-laying rate increases significantly when worms face the anode and the response is EF-dependent, i.e. stronger (6V/cm) and longer EF (40s) exposure result in a shorter egg laying response duration. Worm aging significantly deteriorates the electric egg-laying behaviour with 88% decrease in the egg-count from Day-1 to Day-4 post young-adult stage. Fluorescent imaging of intracellular calcium dynamics in the main parts of the egg-laying neural circuit demonstrate the involvement and sensitivity of the serotonergic hermaphrodite specific neurons (HSNs), vulva muscles, and ventral cord neurons to the EF. HSN mutation also results in a reduced rate of electric egg-laying allowing the use of this technique for cellular screening and mapping of the neural basis of electrosensation in *C. elegans*. This novel assay can be parallelized and performed in a high-throughput manner for drug and gene screening applications.

## 1. INTRODUCTION

A core question in neuroscience is to interrogate how sensory neurons are involved in eliciting a particular behavior. Perceiving a sensory stimulus by the central nervous system induces a flow of electric signals through the motor neurons to stimulate muscle contraction and behavior generation. Due to the complexity of the human brain, model organisms with less complicated nervous system are used to unravel the functions of sensory-motor processes ^[1]^.

The nematode *Caenorhabditis elegans* (*C. elegans*) is a free-living organism that has a simple and well-defined nervous system with a fully sequenced genome which makes it appealing for neuroscience research ^[2,3]^. Although a hermaphrodite worm only possesses 302 neurons, it can perceive and respond to a wide range of stimuli such as chemicals, light, temperature, electric field (EF), and touch through the developed sensory amphid, phasmid, labial, and mechanosensory neurons ^[4–9]^. *C. elegans* behaviors, including movement, egg-laying, and food intake are regulated by these sensory modalities and behavioral screening provides a great means to study neuronal functions and processes in *C. elegans*.

Electrotaxis is one of several behaviors preserved across many species, which is the ability of the organism to sense and move towards a desired direction in the presence of EF ^[10,11]^. Gabel et al. ^[12]^ investigated the neuronal basis of electrotaxis using laser ablation of specific neurons in various transgenic *C. elegans* lines. ASJ and ASH neurons were highlighted as the primary electrotaxis sensory neurons, whereas RIM and AVA were reported to participate in reorientation maneuvers. *C. elegans* electrotaxis was found to be dependent on multiple genes such as *che-2, che-13, eat-4, osm-3, osm-5, osm-6, osm-10*, and *tax-6*. Recently, Chrisman et al. ^[13]^ elucidated the role of the AWC neuron pairs (AWC^OFF^ and AWC^ON^) on electrotaxis and showed that genetic ablation of the neurons is less disruptive than the loss of function asymmetry.

*C. elegans* electrotaxis has also been studied in more controlled microenvironments using microfluidic devices ^[14–19]^. Rezai et al. ^[15,16]^ were the first to exploit microfluidics for the investigation of electrotaxis under direct current (DC) ^[15]^ and alternating current (AC) ^[16]^ EF by measuring the worms’ speed at different larval stages. Since then, electrotaxis has been used in various sorting applications ^[18–20]^ and as a tool for drug screening ^[14,17]^. Most studies focused on the larval and young adult stages, giving less attention to the response at later adult ages. Moreover, investigations were limited to gait behavior in terms of speed, body bend frequency, and reorientation. Herein, we discovered that at later developmental stages, *C. elegans* exhibited unusual and less robust electrotaxis behavior and were interestingly stimulated to lay eggs in a microchannel, a phenomenon that has not been shown to date.

*C. elegans* egg-laying is an established rhythmic behavior of interest to the neuroscience community. It is controlled by a simple neural circuit that can be excited to produce a consistent behavioral output ^[1,21]^. *C. elegans* egg-laying involves interactions between vulva muscles (vms), two hermaphrodite specific neurons (HSN), and six ventral cord (VC) neurons ^[1]^. Worms lay eggs in a temporal stochastic manner that fluctuates between active state, during which the worm lay eggs, and inactive state, during which no egg-laying occurs. Several environmental factors can stimulate or halt egg-laying ^[22]^. For instance, Sawin et al. ^[23]^ reported that mechanical stimulation using vibration inhibits egg-laying. Worms also halt egg-laying in hypertonic salt solutions such as M9 and in the absence of food. Fenk et al. ^[24]^ showed that egg-laying is inhibited by CO_2_ exposure through modulation of the AWC olfactory neurons. Recently, Collins et al. ^[1]^ attempted to understand the egg-laying behavior and the function of each part of the circuit during the active and inactive states. During the active state, the overall behavior was phased with locomotion, while vms were activated by HSN and VC neurons to contract and expel fertilized eggs. Optogenetics is the primary method used to stimulate neurons to study the sensorimotor pathways involved in egg-laying ^[1]^. However, it is restricted to genetically modified worms with light-sensitive ion channels, calling for a simple, on-demand, and inclusive egg-laying stimulation technique applicable to wild-type worms. In this endeavor, we propose the use of DC EF in a microchannel as a stimulus to evoke the egg-laying circuit and induce egg-laying on-demand.

In this paper, we demonstrate for the first time that DC EF evokes the *C. elegans* egg-laying neural circuit. In this context, a simple mono-layer microfluidic device was developed with end-electrodes for electric stimulation and an electrical trap to confine the worm in the field of view while allowing regular egg release. Moreover, the effects of adult worms age and EF direction, strength, and exposure duration on the number of eggs released were investigated. The worms’ muscle activity was quantified through the rate of contraction and relaxation under the influence of EF, and we showed that it was independent of the EF direction. Transgenic lines expressing the fluorescent Ca^2+^ reporter GCaMP5 in HSNs, vms, and VCs neurons and an HSN mutant line were used to investigate the electric egg-laying behavior. Our technique can be used potentially for egg collection and synchronization as well as chemical and mutant screening. Moreover, it will aid in the investigation and mapping of the neural basis of electrosensation in *C. elegans*.

## 2. Materials and Methods

### 2.1. *C. elegans* strains and growth

Nematodes were grown at 20 °C on standard nematode-growth medium (NGM) plates seeded with *Escherichia coli* (*E. coli*) strain OP50 as a food source ^[25]^. OP50 *E. coli* bacteria were cultured in freshly prepared Luria Broth (LB) media (10 g Bacto-tryptone, 5 g Bacto-yeast, and 5 g NaCl in 1 L distilled water) overnight in a thermal shaker incubator at 37 °C. On the following day, 100 μL of the bacterial culture were used to seed the NGM plates. The strains used in this study are showing in Table 1, and they were obtained from the *Caenorhabditis Genetics Center* (University of Minnesota, USA). All experiments were performed under Biosafety Number 02-19 issued by York University’s Biosafety Committee to PR.

**Table 1:**
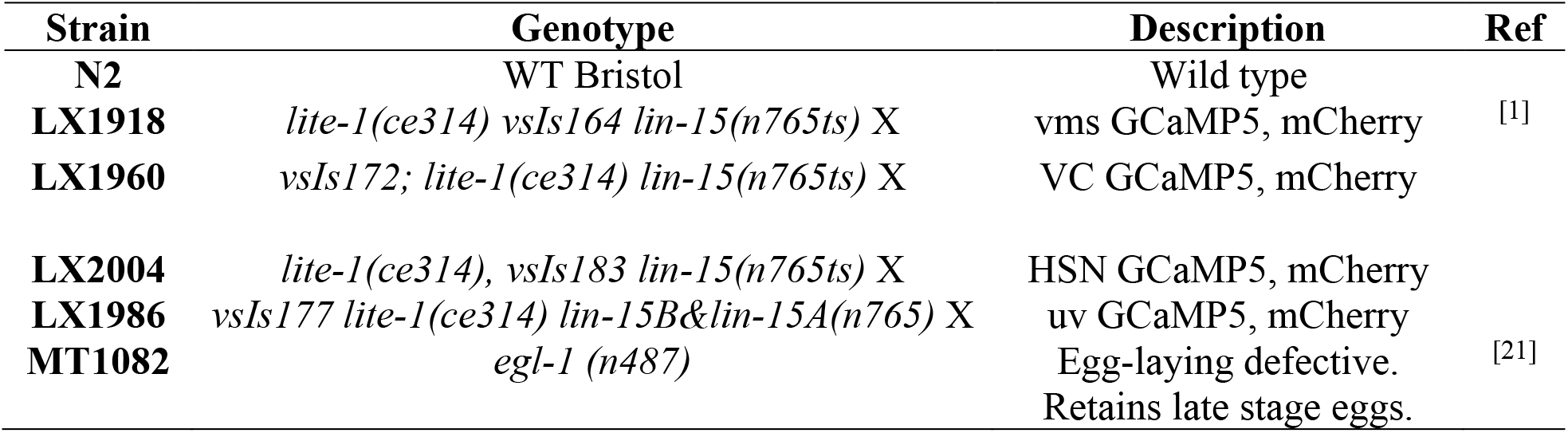
C. elegans strains used in this study.

In all experiments, age-synchronized, well-fed young adult worms were obtained using the Alkaline hypochlorite treatment ^[26]^. Briefly, gravid hermaphrodite worms were collected in a 15 mL Eppendorf tube and treated for 10 minutes with a solution of 3.875 mL double-distilled water, 125 μL NaOH, and 1 mL commercial bleach. Following the treatment, the eggs were obtained by centrifuging the sample at 1500 rpm and incubated at 20 °C overnight in 1 mL M9 buffer (3 g KH2PO4, 6 g Na2HPO4, 5 g NaCl, and 1 ml 1 M MgSO4 in 1 L distilled water) using a RotoFlex™ tube rotator (RK-04397-40, Cole-Parmer, Canada). On the following day, the hatched larvae were plated on freshly prepared NGM plates to allow for standard growth. All experiments were conducted using worms at day one post young adult stage (~64 hr or D1), except for age effect experiments in which the worms were transferred to a new NGM plate for consecutive day studies in D2 to D4.

### 2.2. Microfluidic chip design and fabrication

The proposed microfluidic device shown in Figure 1A was designed to study the effect of DC EF on the egg-laying behaviour of *C. elegans* and to provide the ability to image the worms fluorescently. The mono-layer microfluidic device consisted of a 3 cm long straight microfluidic channel (300 μm wide) with a 1.3 mm long mid-section electrical trap (100 μm wide) to provide a symmetrical condition for the EF. All channels were 65 μm thick. The two end reservoirs were connected to two copper-wire electrodes. The electric trap enlarged in Figure 1A was used as the test area to confine the worm’s movement for studying the effect of EF direction, magnitude and duration, as well as worm age on EF induced egg-laying. The confinement was required to lessen the worm movement for easy monitoring of egg laying and fluorescent imaging. Moreover, numerical modeling was conducted to confirm the uniformity in the EF distribution along the channel when EF was applied (supplementary file section 1).

**Figure 1:**
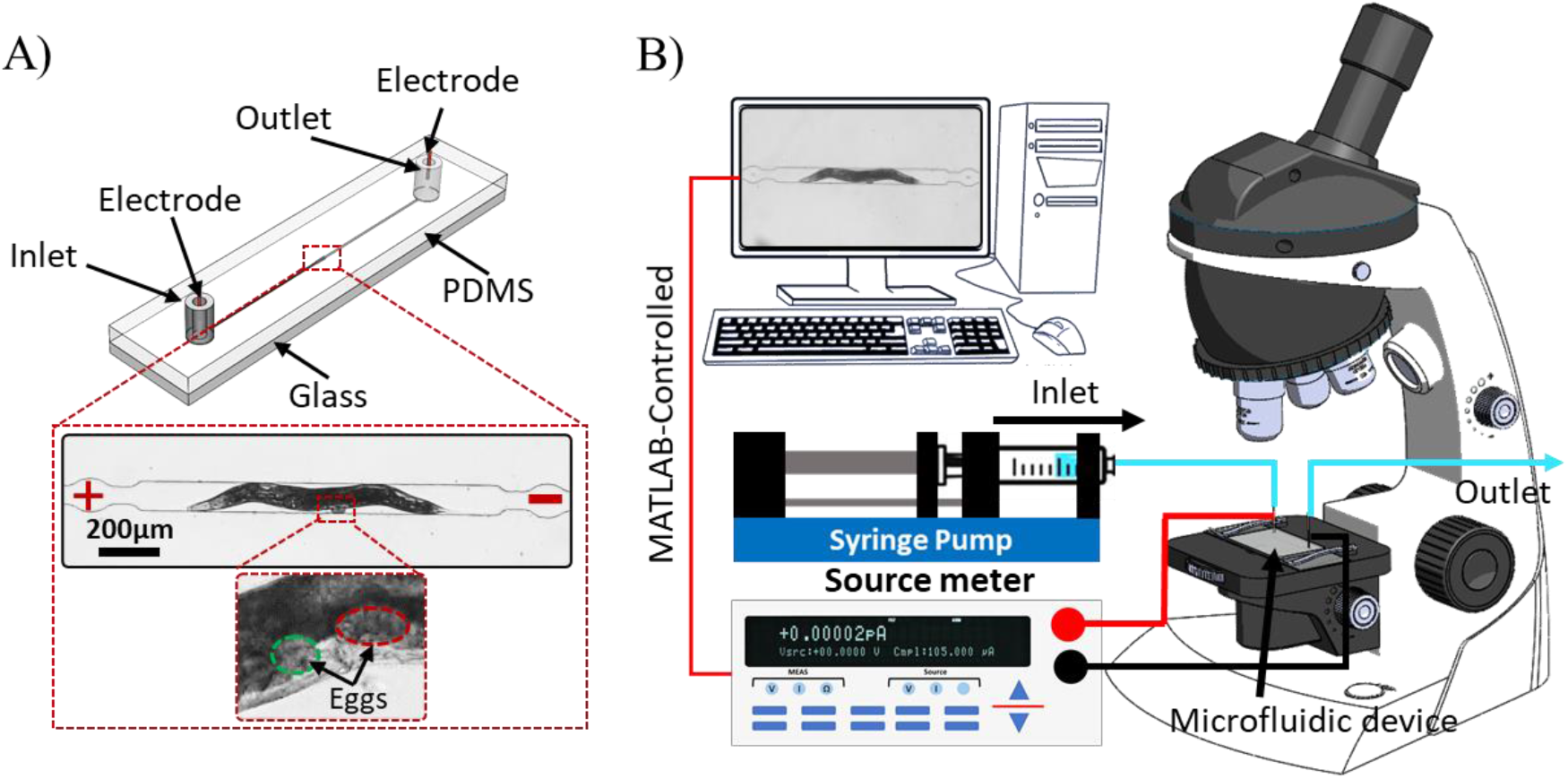
(A) Microfluidic device (2.5cm×1cm) consisting of an inlet, an outlet, and a symmetrical microchannel with a mid-trap for worm testing and two end electrodes for EF stimulation. (B) Experimental setup consisting of our microfluidic device, a microscope, a syringe pump, and an electrical sourcemeter.

The microfluidic device was fabricated using standard photo- and soft-lithography techniques ^[27]^. To fabricate a negative replica, a 65 μm thick layer of SU-8 2075 photoresist (MicroChem Corporation, USA) was photolithographically patterned on a 4-inch diameter and 500 μm thick silicon wafer (Wafer World Inc., USA) using UV exposure at 365 nm with 11.1 mW/cm^2^ (UV-KUB 2, KLOE, France) for 14s. Inlet and outlet Masterflex tubes (L/S 14 size, Gelsenkirchen, Germany) were placed over the master mold. Then, polydimethylsiloxane (PDMS, Dow Corning, USA) elastomer and curing agent were mixed at 10:1 ratio and poured over the master mold followed by curing for 2 hours at 80°C. Then, the cured PDMS layer was peeled off the master mold, bonded to a pre-cleaned glass substrate (75×25 mm^2^) by oxygen plasma (PDC-001-HP Harrick Plasma, USA) at 870 mTorr pressure and 30 W power for 30 s, and kept for 30 min at 65 °C to enhance bonding. EF stimulation was attained by punching two 3 cm long copper electrodes through the PDMS into the inlet and outlet tubes. Liquid PDMS mixture was used to seal the surrounding areas of the copper insertions to prevent leakage.

### 2.3. Experimental setup and procedures

The microfluidic device was installed on an inverted microscope (DMIL LED Inverted Routine Fluorescence Microscope, Leica, Germany), equipped with a color camera (MC170 HD, Leica, Germany), as shown in Figure 1B. The two copper electrodes from the device were connected to a DC sourcemeter (Model 2410, Keithley Instruments Inc., USA) for EF stimulation. The sourcemeter was controlled using a custom-developed MATLAB code to regulate the applied voltage, stimulation time, and EF direction. The EF in the trap was varied from 2 to 8 V/cm, and the stimulation time was varied from 1 to 40 s. This range of EF in the trap was determined by a simple COMSOL Multiphysics model, after applying a voltage of 2.25 to 9 V across the channel, as described in supplementary file section 1. EF direction was decided based on the worm’s head orientation in the channel after loading, i.e. anode or cathode stated when the worm head was towards the positive or negative electrodes, respectively.

Prior to the experiments, the device was filled with M9 buffer using a manual syringe connected to the inlet. The outlet was connected using a valve to two 15 mL centrifuge tubes, one for device washing and the other for egg-collection when needed. A single worm from a synchronized population (D1 to D4 post young adult) was manually picked (Wormstuff, USA) and loaded into the inlet tube of the device. Then, a 10 mL syringe filled with M9 was used to guide the worm to the middle electric trap manually and, once the worm reached the trap, the flow was stopped, and the worm was left for 60 s to acclimate to the environment before exposure to the EF. Next, the MATLAB code was run, and a series of step voltage pulses were initiated based on the desired EF and exposure duration. The EF pulse had an ON state of 1 to 40 s and an OFF state of 25 s for a fixed total exposure time of 10 min for each worm. Results were recorded in terms of the number of eggs and the time of egg-laying for behavioral assay. Video recording under fluorescent setting was done during neuron and muscle transient response assays. Then, the worm was flushed out of the chip to allow for the next worm to be tested.

### 2.4. Video processing and phenotypic analysis

The camera mounted on the microscope was used for acquiring movie clips of the experiments at 5x magnification with 30 frames per second (FPS) speed and 5 megapixel resolution. Our custom-written MATLAB code was used to process the recorded videos and detect the worms’ centerline. For this, the user was asked whether to perform worm length measurement or fluorescent intensity analysis. If length analysis was selected, the user was asked to import a video and the first frame of the imported video was shown on the screen (Figure 2Ai). The user was then asked to import the background image (Figure 2Aii) which was taken from the chip after unloading the worm. The background image was subtracted from each video frame (Figure 2Aiii) and binarized (Figure 2Aiv), achieving high-quality binarization of the worm image with all the other objects excluded from the image. For instance, Figure 2Aiii shows an egg that was removed from the binary image in Figure 2Aiv. Then, the white pixels in Figure 2Aiv were used to obtain the centerline of the worm. The centerline was determined by rearranging the white pixels into verticale lines and getting the middle pixel of each vertical line in the X-direction. By connecting these pixels, the centerline (Figure 2Av) was determined and used to measure the worms’ length.

**Figure 2:**
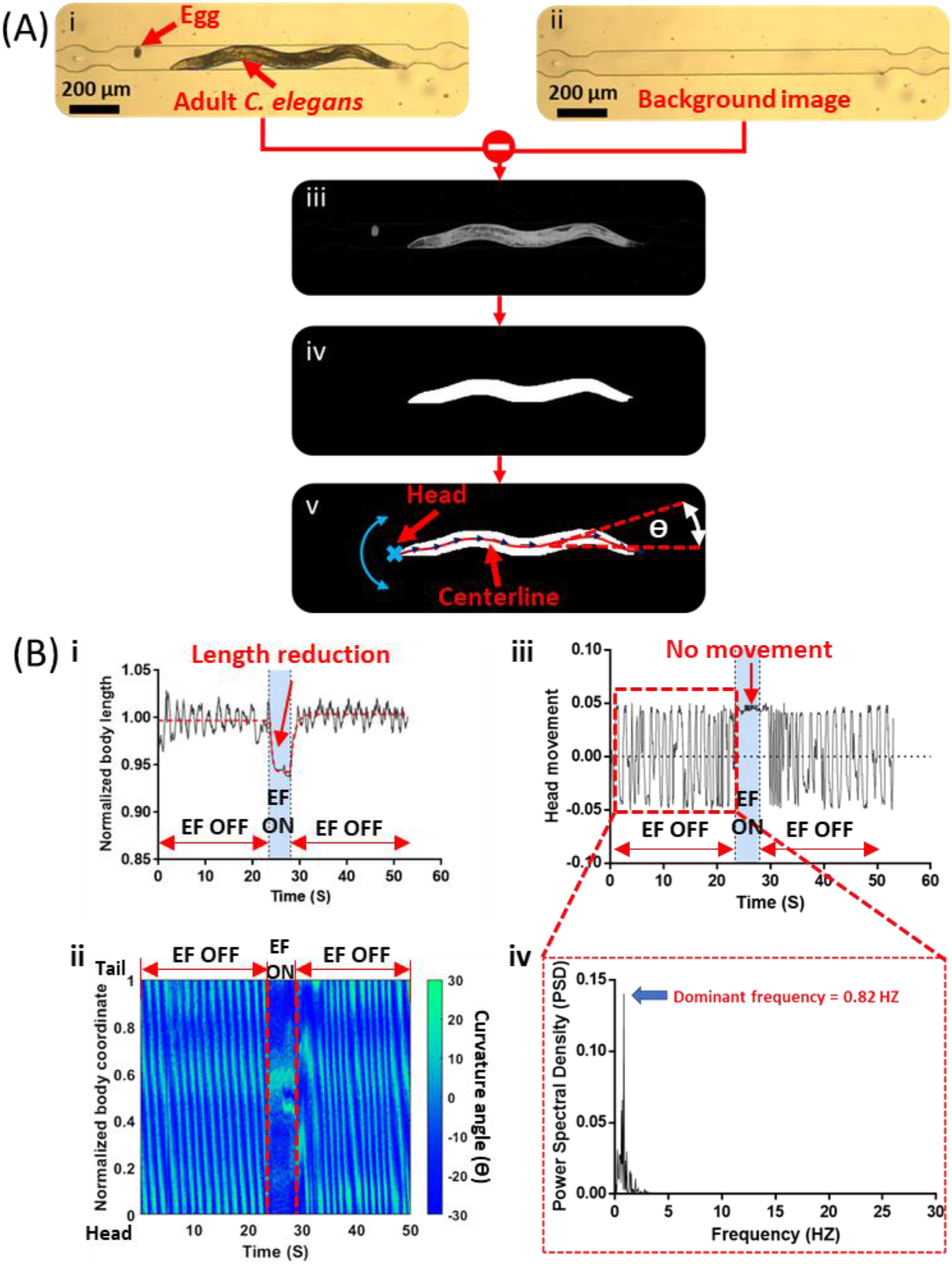
Details of the MATLAB-based image processing code for frame by frame analysis of the recorded videos. (A) (i) A D1 adult worm captured inside the electric trap. (ii) The background image after pushing the worm out of the trap. (iii) Difference between (i) and (ii) to obtain an image with only the worm and the eggs. (iv) Binarized image and the largest connected object (the worm) retained. (v) The worm’s centerline, head location and body segment curvature angle used for analysis. (B) (i) Normalized body length, (ii) body curvature plot, and (iii) head movement tracking of the worm inside the microfluidic device. Head tracking data was used to determine the dominant frequency in (iv) by taking the power spectrum density.

The worm’s length was normalized by its length before EF exposure and plotted over time (Figure 2Bi), then used as a readout for the electrically-induced body muscle contraction and relaxation. For these experiments, six worms were analyzed for three consecutive cycles of anode or cathode stimulation, giving a total of 18 measurements. Each cycle involved an excitation period of 5 s preceded and followed by 25 s acclimation periods. As shown in Figure 2Bi, during the first 25 s, the worm length was not significantly changing; however, when the EF was turned on, the worm’s length contracted for 5s with an exponential-like decay. Then, when the EF was turned off, the worm started to relax, and the length returned to its original value rapidly (Supplementary Video S1).

The worms’ body movement was determined by tracking the body sinusoidal wave with time to derive the body curvature plots in Figure 2Bii. These plots characterized how the bending waves propagated in time along the worm body, which were used to determine the swimming frequency during the on- and off-EF periods (Supplementary Video S1). Worm body curvature was represented in terms of the tangent angle made by each body segment (θ in Figure 2Av). Accordingly, Figure 2Bii depicts the curvature angle θ of different worm body segments (0 for the head and 1 for the tail) as a function of time. A vertical line across the graph in Figure 2Bii represents the sinusoidal shape of the worm at a specific time frame, whereas, the observer can track the movement of specific body segments as a function of time by sketching a horizontal line. Moreover, the worm’s head, symbolized by a cross sign in Figure 2Av, was monitored to quanitfy the movement frequency by tracking the lateral head movement with time (Figure 2Biii and 2Biv). For instance, Figure 2Biii shows a pattern of continuous sinusoidal waves during the EF-Off period, followed by a pause period when the EF was applied and a resumed sinusoidal head movement pattern in the post-exposure period. Figure 2Biv shows the power sepctral destiny analysis to determine the dominant frequency for the siusoidal head movement which was obtained by our MATLAB code.

Fluorescent intensity analysis was also done in our MATLAB code for the GCAMP strains reported in Table 1. Before applying the EF, it was necessary to ensure that the neurons or muscles reached the baseline activity before recording the video. Once the video was imported, the user was asked to select a region of interest (ROI), for instance around the vulva. In each video frame, the mean green flourescent intenisty of the ROI was determined (I_ROI_), subtracted from the background intensity (F = I_ROI_- I_background_), normalized to the mean baseline activity 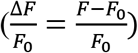, and plotted versus time. The baseline activity (F_0_) was quantified as the mean green fourescent intensity of the ROI prior to the stimulus. The background intenisty was obtained through selecting another ROI, similar in size to the target ROI, within the sides of the frame.

### 2.5. Off-chip locomotion assays

For off-chip locomotion analysis, the electrically exposed worms were collected after the experiment and transferred into 3 cm diameter Petri dishes containing 1 ml of M9 buffer. An upright Leica stereomicroscope (Leica MZ10 F fluorescence microscope, Leica, Germany) was used to video-record the worms swimming behaviour in M9. Then, the videos were post-processed using the worm tracker plugin in Fiji software^[28]^ to subtract the background and determine the average speed and swimming frequency. For comparison purposes, worms from the same population and age were transferred into M9, without exposure to EF, and their speed and swimming frequency were recorded as the control group (supplementary file section 2).

### 2.6. Statistical analysis

A pilot experiment was conducted to provide a base for determining the sample size using GPower software ^[29]^. At least two replicates of 10 worms were used in the egg-laying experiments, and at least five worms were used for the length reduction, body bend, and fluorescent intensity analysis. The exact sample size used for each experiment is presented in the figure captions. The data were presented in two formats, either as the mean ± standard error of the mean (SEM) or using box plots with medians, 25% and 75% percentiles, and maximum and minimum data points. We used the Mann-Whitney test to determine the statistical significance between two groups, while the statistical differences among multiple data points were determined using one-way ANOVA analysis. The significance levels were identified by stars, i.e. * for p-value<0.05, ** for p-value<0.01, *** for p-value<0.001, and **** for p-value<0.0001.

## 3. RESULTS AND DISCUSSION

### 3.1. Electric egg-laying of C. elegans

In a DC EF, *C. elegans* sense the strength and direction of the EF with amphid neurons and swim towards the negative pole on open agarose surfaces ^[12]^ and inside microchannels ^[15]^. Aging has been shown to decrease the electrotactic response by 70% from young adult to day-8 adult stage ^[30]^. Here, we asked for the first time whether EF induces any behavior other than electrotaxis in *C. elegans*, especially at gravid adult stages that electrotaxis weakens significantly. To answer this question, two microfluidic devices, one with a 300μm-wide channel and one with a 100μm-wide trap in the middle as shown in Figure 1A, were used. The experimental setup demonstrated in Figure 1B was used for electric stimulation of the worms as elaborately discussed in the Materials and Methods section. Video processing and analysis was done to assess various movement phenotypes that are shown in Figure 2 and discussed in the Materials and Methods section.

We fabricated a 4cm-long and 300μm-wide microfluidic electrotaxis channel ^[15]^ and tested the effect of EF on gravid adult worms. In these preliminary assays, worms were first facing towards the cathode, however when the EF was reversed, they showed a tendency to turn slowly towards the cathode and while doing so, they also deposited eggs ((Supplementary Video S2)). For example, Figure 3A shows sequential images of a D1 adult worm exposed to a 6V/cm EF, showing egg-laying behaviour instead of electrotaxis over a period of 5s. Counting the number of eggs and neuronal imaging of the worm in this wide-channel device imposed some challenges mainly due to the continuous and wide-range movement of the worm during egg deposition.

**Figure 3:**
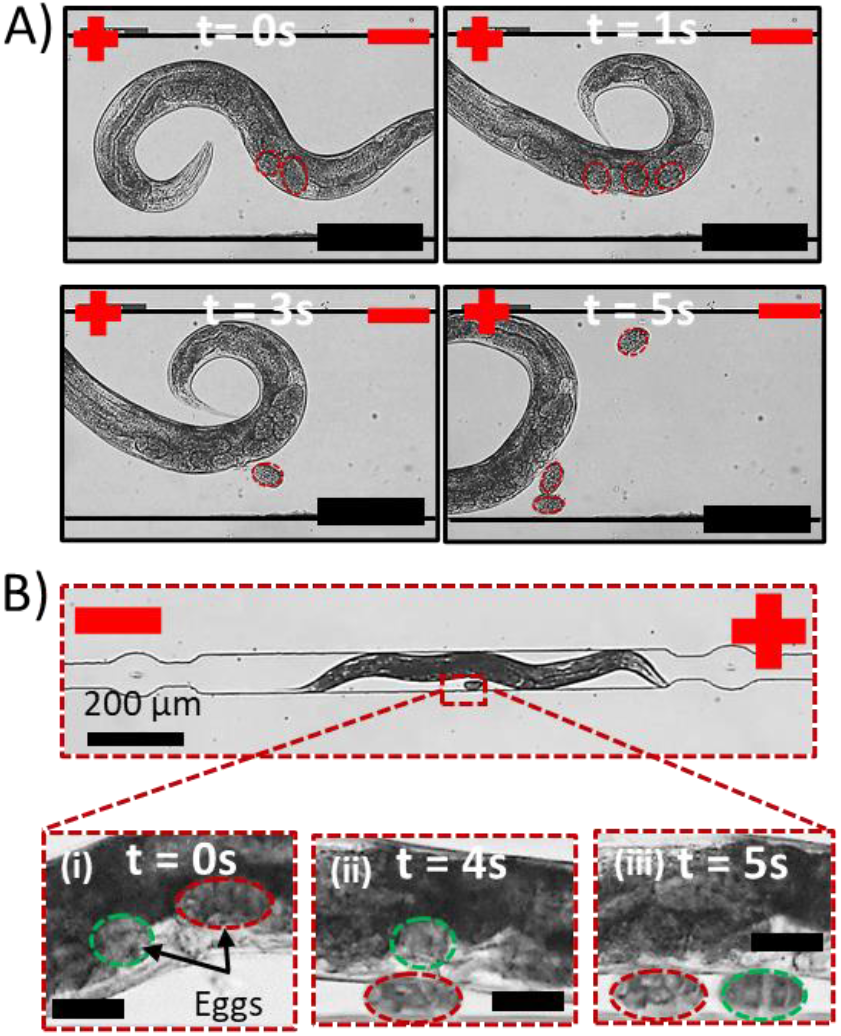
(A) Electrotaxis re-orientation attempt of a D1 C. elegans in response to an EF of 6 V/cm which was accompanied by egg-laying inside a 300 μm wide microchannel. (B) Electric egg-laying behaviour of a D1 C. elegans in response to an EF of 6 V/cm inside the 100 μm wide electric trap of the proposed microchannel.

To address the limitations above, the microfluidic device shown in Figure 1 was developed to restrain the worms’ orientation while allowing egg release and fluorescent imaging during EF exposure. Figure 3B shows sequential images of a D1 adult *C. elegans* in the electric trap of our device while being exposed to an EF of 6 V/cm with the cathode positioned at the tail of the worm. By applying the EF, we observed that although the worm could not rotate towards the cathode, it still deposited eggs over a period of 5 s. We also noticed that the egg-laying phenomena was accompanied by various movement kinematics in this device, including body shortening and immobilization which aided in on-chip neuron imaging of GCaMP strains as discussed later in this paper.

The discovery of *C. elegans* EF induced egg-laying in a close-fitting microchannel led us towards in-depth investigation of the effect of electric pulse direction, strength, and duration as well as worm age on the egg-laying behaviour in the following sections. We have also examined the involvement of neurons and muscles in electric egg laying.

### 3.2. Effect of EF direction on the egg-laying behaviour

D1 adult worms (N=70) were positioned in the electrical trap of the microfluidic device and exposed to EF with cathode at their heads or tails. In the absence of EF, the worms moved in the trap with an average head movement frequency of 0.89 ± 0.18 Hz and no significant change in their swimming kinematics (Supplementary Video S1). Moreover, they showed no egg-laying behaviour during a maximum waiting time of 10 min. Then, a series of 6 V/cm EF pulses (5s on and 25 s off) was applied for 10 minutes while the worms were facing either the cathode or the anode. Supplementary Video 3 demonstrates the behaviour of one of these worms before, during and after exposure to EF with cathode at its tail. As shown in Figure 4A, anode-facing worms deposited significantly more eggs in response to the EF (9 eggs/worm in average) compared to the cathode-facing worms (1 egg/worm) (Supplementary Video S3). It is worth mentioning that from the 70 worms tested in each group, the egg-laying response rate was 92.8% versus 14.3% for the anode- and cathode-facing worms, respectively. Interestingly, this robustness in egg-laying response for the anode-facing worms is similar to *C. elegans* electrotaxis response rate when cathode is positioned at the posterior side of the worms.

**Figure 4:**
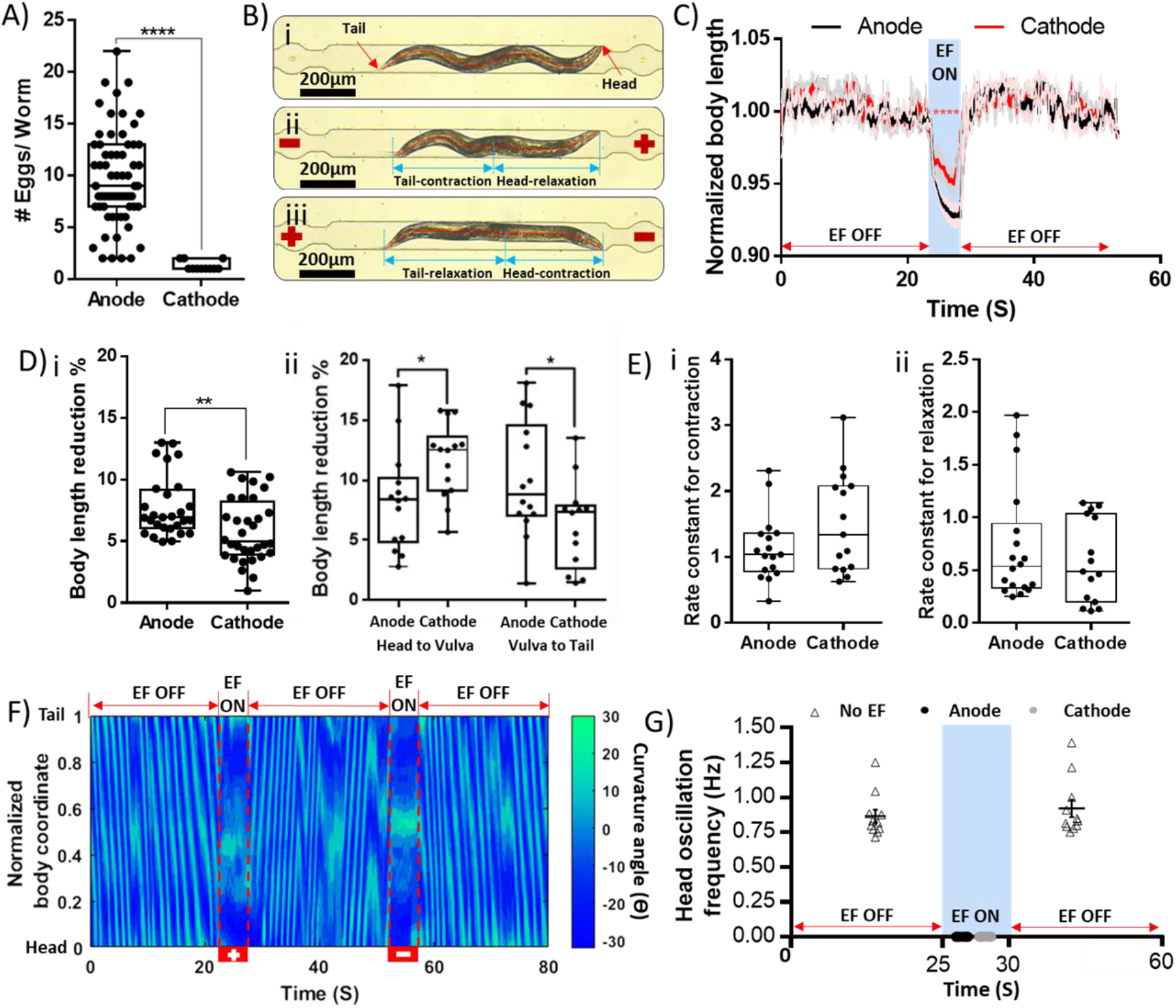
The effect of EF direction on C. elegans egg-laying behaviour. EF of 6 V/cm was applied for up to 10 min in pulses with 5s on and 25s off cycles. (A) Number of eggs per worm for N = 70 worms that were facing towards the anode or cathode in the trap. (B) Optical microscopic images showing the worm in the electric trap during (i) no EF, (ii) anode-facing EF, (iii) cathode-facing EF. (C) Instantenous normalized body length of anode-and cathode-facing worms (N=18), showing contraction and relaxation periods fitted with one-phase decay and association models, respectively, to quantify the contraction-relaxation kinematics. (D) (i) Maximum total body length reduction and (ii) head-to-vulva and vulva-to-tail length reduction of worms facing either the anode or the cathode. (E) Rate constants for (i) contraction and (ii) relaxation. (F) Time-lapse normalized body curvature angle of a worm exposed to two EF pulses with anode and then cathode at head. (G) Worms head oscillation frequency during the on- and off-periods of the EF, showing a complete movement inhibition during EF stimulation for both anode-and cathode-facing worms.

To investigate the causes and effects behind the EF-induced egg-deposition, various locomotion kinematics such as changes in the worms’ body length (Figure 4B) and the head movement frequency were studied parametrically before, during and after EF exposure. Figure 4C depicts the instantaneous length reduction due to EF exposure for anode- and cathode-facing worms. A significant shortening in the worms’ length was observed in both cases, implying involvement of the body wall muscles in egg-deposition during EF exposure. However, the anode-facing worms had an average length contraction of 7.75%, which was significantly (p<0.01) larger than the 5.8% contraction of the cathode-facing worms, as shown in Figure 4Di. This corresponds well with the larger number of eggs released in the worms initially facing the anode as shown in Figure 4A.

During the experiments, we also noticed that the muscle contraction-relaxation behaviour was not consistent between the anode- and cathode-facing cases. For instance, in the anode-oriented worms, we observed a dominant contraction in the mid-region body muscles (Figure 4Bii) while for the cathode-oriented worms, the contraction was significant in the headwall muscles as shown in Figure 4Biii. To depict this quantitatively, body length reduction was analyzed separately on the head-to-vulva and vulva-to-tail sections of the worms, before and during EF exposure in anode- and cathode-facing worms (Figure 4Dii). The results confirmed that contraction was associated with the head region in the cathode-facing worms while the anode-facing worms experienced dominant contraction in the mid-body or the tail region. In other words, regardless of orientation, the side of the worm facing the cathode experienced a more dominant contraction of the body. Moreover, contraction in the mid-body region where the vulva is located may be a reason for the worms expelling more eggs when facing the anode, i.e. exerting more force on the vulva muscles to twitch and lay eggs.

Muscle excitation-contraction coupling is an interesting process by which *C. elegans* crawl on an agar surface or swim in liquids. Having seen the effect of EF on muscles in our experiments, we became interested in examining the contraction-relaxation characteristics of the muscles. The data in Figure 4C were further analyzed and fitted using an exponential decay or growth equation to determine how fast muscles contract or relax during and after EF activation. We aimed to figure out whether there is a difference in the contraction/relaxation kinetics under the two directional EF stimulations. Figure 4E shows the rate constant of relaxation and contraction for anode- and cathode excitations. The results indicated no significant difference in the contraction and relaxation rates of both experimental conditions. Moreover, our data matched the contraction and relaxation rates reported by optogenetics for unexposed worms ^[31]^. This showed the potential use of our technique in studying the overall muscle kinetics of wild-type and potentially other worm strains.

Another interesting parameter observed in both anode- and cathode-facing worms was the loss of the swimming kinematics during the EF exposure, determined by the worms’ body movement angle and head movement frequency. The body curvature of the worm in Figure 4F fluctuated between ± 30 degrees during the off-EF period, whereas it became constant during EF exposure due to the worm ceasing its movement as shown in Figure 4G. The calculated swimming frequency during the off-EF period was 0.87± 0.046 Hz and 0.92± 0.06 Hz before and after the stimulus, respectively, whereas, during the on-EF period, the frequency reduced to zero. This further confirmed that during EF exposure, the worms ceased movement, contracted their bodies, and deposited eggs in the device.

### 3.3. Effect of EF strength on the egg-laying behaviour

Having established the importance of worms orientation towards the anode for electric egg-laying, here, we asked if EF strength has a dominant effect on this novel behaviour. Figure 5A shows the effect of 5s-long EF pulses with 2-8 V/cm strengths on the egg-laying, body length and oscillation frequency of D1 wildtype worms facing the anode. The results showed a robust and peaking egg-laying activity at EF=6 V/cm while a 2 V/cm EF did not induce any egg-laying (therefore omitted from Figure 5). At low EF of 4 V/cm, a significant decrease in egg-laying was observed compared to 6 and 8 V/cm. Although an EF of 8 V/cm stimulated the worms to expel eggs, the number of extracted eggs significantly decreased compared to 6 V/cm.

**Figure 5:**
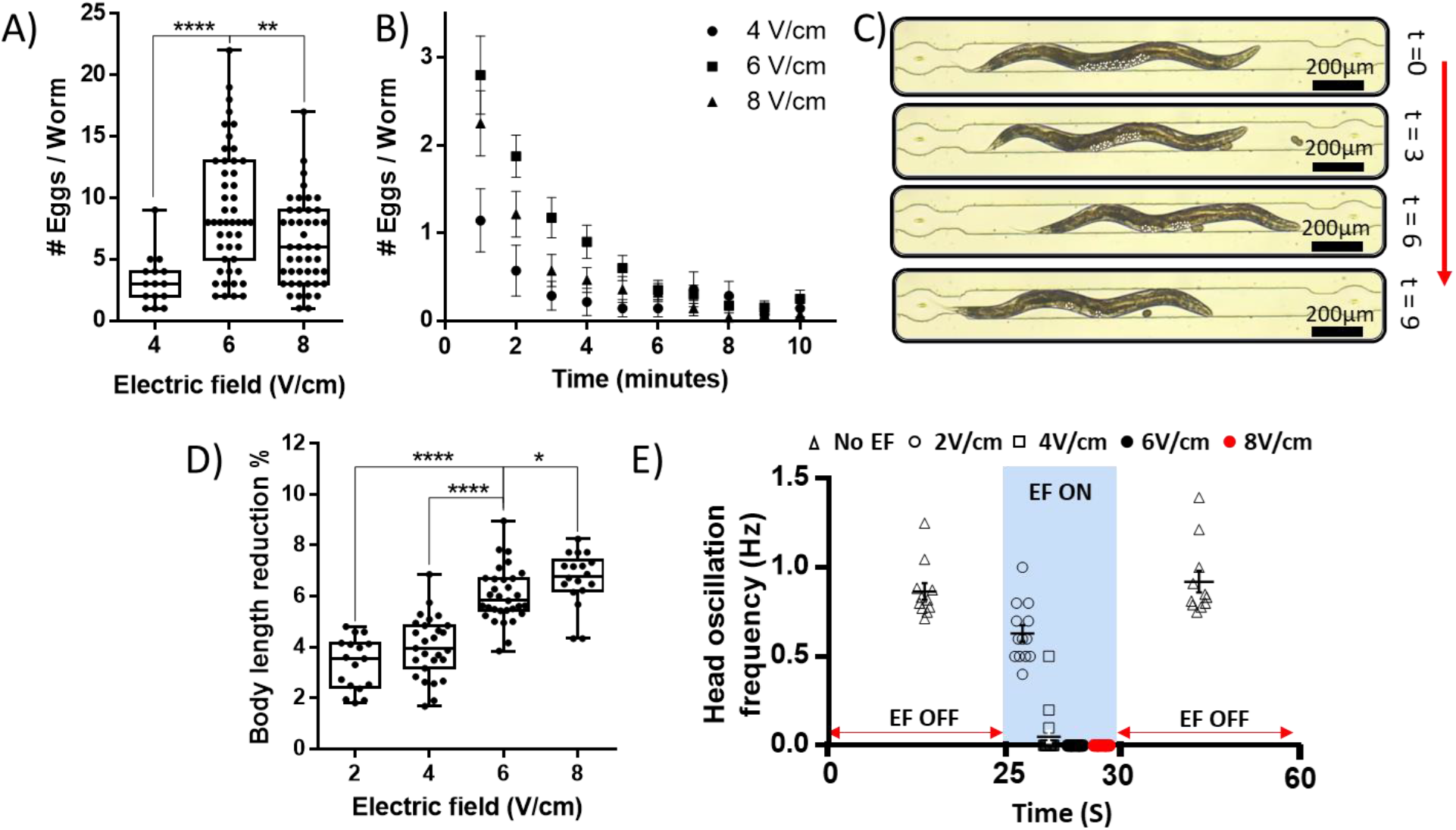
The effect of EF strength on egg-laying behaviour of D1 anode-facing wildtype worms using 5s on and 25s off pulses for 10 min. Number of eggs per worm for at least N=20 worms at different EF strengths is shown (A) for the whole 10 min period and (B) at the end of each minute. (C) Time lapse images showing egg-laying and emptying the uterus without reproducing new eggs (time is represented in minutes). (D) Relative body length reduction percentages during EF stimulation at different EFs. (E) Worm head movement frequency during on- and off-periods of EF at different EF strengths.

To better understand the effect of EF strength, we analyzed the rate of egg-laying per minute over the 10 min stimulation period (Figure 5B). At all times during the experiment, the number of eggs expelled at 6 V/cm was more than 4 and 8 V/cm EFs. Egg-release rate decreased with time for all EFs and reached a minimum value at approximately 6 min post EF exposure, after which the rate plateaued. This was because *C. elegans* hermaphrodites usually store around 10-15 fertilized eggs in their uterus. Naturally, the worms give birth to 2-3 eggs every 20-25 min ^[21]^, which means that their capability to sustain a constant number of eggs in the uterus is limited under normal conditions. In our microfluidic chip, most eggs were expelled in the first 4 min, and the worms were not able to produce new eggs during the 10 min experimental period. This was confirmed by time-lapse optical images of worms during the 10 min period in Figure 5C, illustrating that the number of eggs in the uterus decreases over time with no new eggs produced.

The effect of EF strength on egg laying may be understood further by performing body length analysis as shown in Figure 5D. Increasing the EF strength consistently resulted in more significant body length shortening, which might be a reason behind expelling more eggs. The shortening effect was most significant when increasing the EF from 4 to 6 V/cm which corresponded well with the significant increase in the rate of egg-laying at this condition. Moreover, it is well understood that *C. elegans* electrotaxis increases with EF in the range of 2-6 V/cm^[15]^, which is the reason behind the worms’ increased tendency to turn towards the cathode in our device, while the channel’s low width was preventing this reaction. Thus, the worms shrank instead of turning and, in response to the shrinkage, they expelled more eggs. The shortening phenotype has also been correlated with the activation of the VC neurons or the body wall muscles ^[1]^ which will be discussed later in this paper. Moreover, the increase in body shortening resulted in a significant decrease in the head movement frequency when increasing the EF from 2 to 8 V/cm as shown in Figure 5E. Lastly, the slight reduction in egg laying at 8 V/cm may be attributed to the EF-induced paralysis which was also reported for freely moving worms by Rezai et al.^[15]^ in wide electrotaxis channels.

### 3.4. Effect of EF pulse duration on the egg-laying behaviour

The research hypothesis investigated in this section was the effect of pulse duration on electric egg laying of adult *C. elegans*. Moreover, extracting more eggs in a short period is of interest to many *C. elegans* laboratories for age-synchronizing the worms. Thus, the exposure time to the electric pulse was varied from 1 to 40 s to investigate its effect on the egg-laying process. Figure 6A shows the total number of eggs released over 10 min by D1 wildtype worms which gradually increased with pulse duration until it reached a constant rate beyond the 5s stimulation period. Figure 6B shows that increasing the pulse duration from 5s to 40s stimulated the worms to expel their eggs significantly faster (Supplementary Video S4), as interpreted by shorter egg-laying durations. The egg-laying response duration was recorded based on when the worm stopped to give an egg for three consecutive pulses at each pulse duration. To better understand the EF pulse effect, we also plotted the time-lapse egg-laying rate for all pulse durations, as shown in Figure 6C. This presentation confirmed that at longer pulses, the worms give more eggs in a shorter overall duration, perhaps due to contractions and vulva muscle openings lingering for longer periods of times during each pulse.

**Figure 6:**
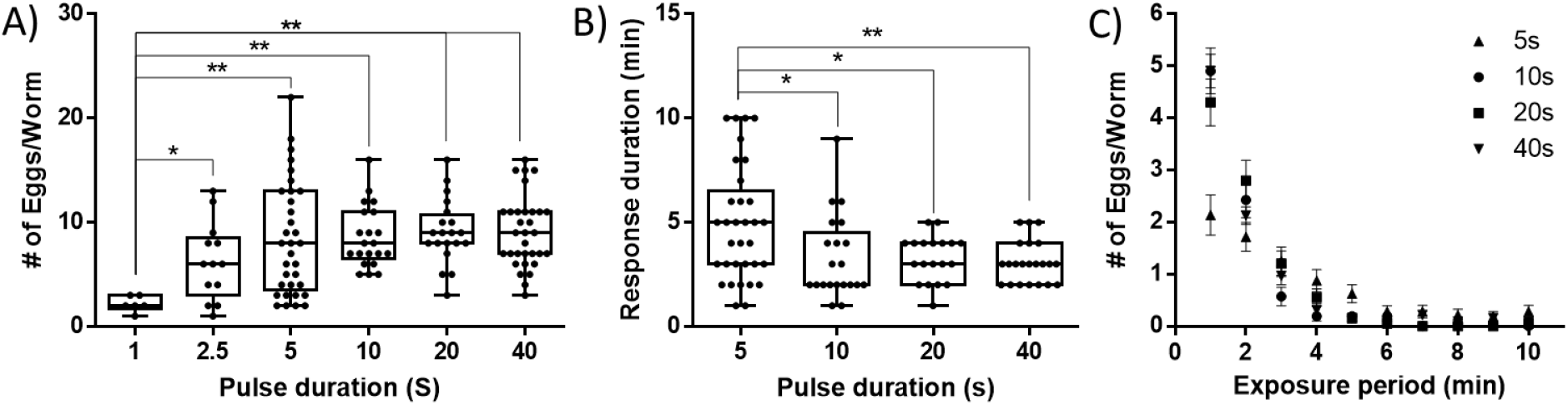
The effect of EF pulse duration on the egg-laying behaviour of D1 anode-facing wildtype worms at 6 V/cm EF. (A) Number of eggs per worm for at least N = 20 worms at different pulse durations. (B) Overall egg-laying response duration at different pulse durations. (C) Time-lapse rate of egg-laying at different pulse durations (legend).

### 3.5. Effect of worm age on the egg-laying behaviour

*C. elegans* has been proposed as a promising model for aging studies.^[32]^ It is well understood that aging causes a progressive decrease in the egg-production rate^[33]^ and electrotaxis of *C. elegans* ^[30]^. However, the effect of aging on electric egg-laying of *C. elegans* has not been studied. Here, we studied the electric egg-laying of worms aged from D1 to D4 past young adult stage using 5s long EF pulses of 6 V/cm for 10 min. Figure 7A shows that worms of all ages were responsive to EF, with a continuous significant decrease in their egg laying from D1 to D4. To compare our data with the typical egg-release results for natural egg-laying (uninduced), we performed an off-chip egg-counting assay for 20 worms at ages D1 to D4, raised on NGM plates. The results in Figure 7B showed a slight decrease in the egg-laying rate from D1 to D2, followed by a significant decline from D2 to D3 and D4. The electric egg-laying results follow the same trend of natural egg-laying, noting that the difference between the number of eggs obtained in Figures 7A (electric) and 7B (natural) is due to differences between the assay times (10 min for electrical and full day for natural). Moreover, the decrease in D4 worm’s response in both cases can be attributed to the age-related deficiency of egg-laying reproductive system, i.e. most of the expelled eggs were lysed or broken once ejected by the vulva.

**Figure 7:**
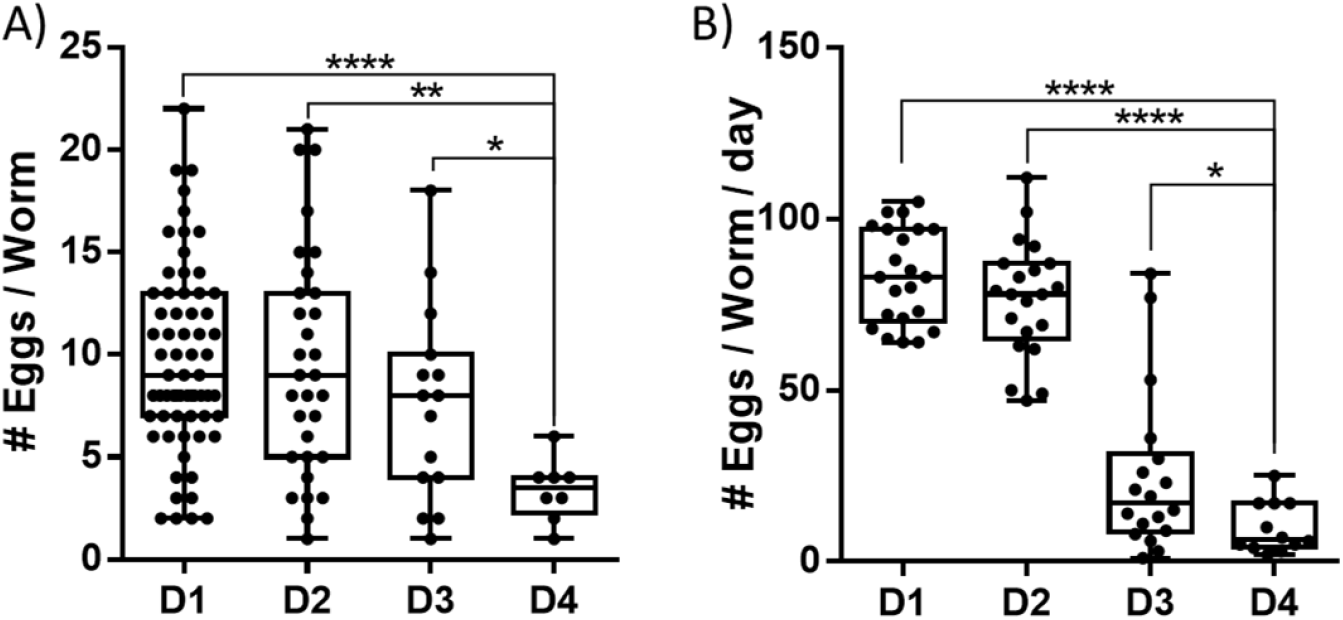
Aging effect on the egg-laying behaviour of (A) anode-facing adult worms at 6V/cm EF and 5 s pulse duration in the microfluidic device, compared with (B) worms natural (uninduced) egg-laying off-Chip on NGM plates.

### 3.6. Neuron and muscle activities during electric egg-laying behaviour

According to *C. elegans* egg-laying wiring diagram in Figure 8A^[1]^, direct synaptic connections between HSN and VC neurons are required to command vms to enter the active state and stimulate egg-laying. Previous studies showed the level of coordination between the activation of the egg-laying neural circuit and egg production.^[1]^ Here, we aimed at investigating the level of coordination between the EF stimulation and the egg-laying neural circuit using the genetically encoded calcium indicator, GCaMP. We chose the transgenic strains LX1918, LX2004, LX1960, and LX1986 (Table 1) because they express the fluorescent Ca^2+^ reporter GCaMP in vms, HSN, VCs neurons, and uv1 cells, respectively. Primarily, we investigated the electric egg-laying of all the strains and observed that they deposit significantly more eggs while facing the anode in response to EF pulses of 6V/cm, which was statistically similar in counts to that of the wildtype worms (Figure 8B).

**Figure 8:**
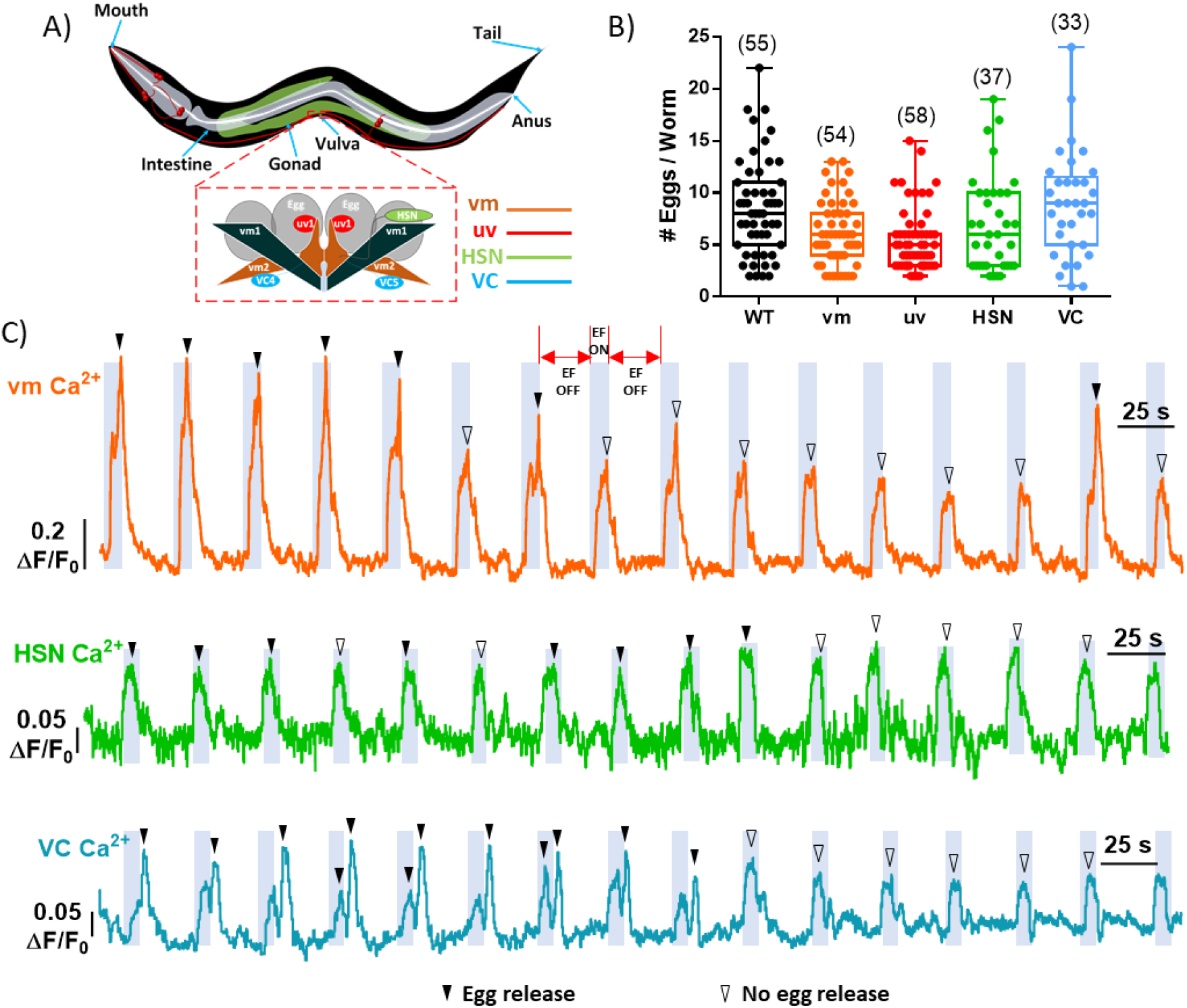
Effect of EF pulses (6 V/cm, 5s on, 25s off) on egg laying neurons and muscles of C. elegans. (A) Schematic of the egg-laying neuromuscular circuit, showing the vms (orange), HSN (green), VC neurons (blue), and uv1 cells (red). (B) Number of eggs per worm for worms facing towards the anode and exposed to EF pulses of 10 minutes. Sample sizes are mentioned between brackets in the graph. (C) Eight-minutes recordings of fluorescent activity in vms, HSN and VC neurons, during the ON (gray-shaded) and OFF (white-shaded) EF periods. Filled arrow heads represent egg-laying events and empty arrow heads represent no-egg laying event.

Using long-term fluorescent imaging of calcium transients, we investigated the activation of the vms, HSNs and VC neurons during electric egg laying in our microfluidic device (Supplementary Video S5-S10). An EF of 6 V/cm was applied as a series of pulses of 5s on and 25s off, while the worms were facing the anode, resulting in 16 consecutive cycles as shown in Figure 8C for representative worms. In all strains, we observed a striking increase in the activity of cells (Figure 8C) due to cell depolarization during the anodal stimulation that was mostly associated with egg-laying events (black filled triangles). No egg release (black empty triangles) happened mostly at the second half of the experiments during pulses 8-16, expectedly due to the depletion of uterus and lack of eggs in the vicinity of the vulva to be released upon pulsation.

In Figure 8C, we were able to distinguish between the strong depolarizations associated with strong twitches of the vms during egg-laying events (black filled triangles) and the weak contractions with no egg-laying events (black empty triangle) (Supplementary Video S5). The same trend was observed for the VC neurons with higher Ca^2+^ transients during egg-laying (Supplementary Video S7). The magnitudes of HSNs activities were not correlated as strongly with egg-release, but all egg-laying events were accompanied by HSN firing (Supplementary Video S9). uv1 Ca^2+^ transients were weak to be recorded using our microscopy thus it has been excluded from the analysis. It is also worth mentioning that we rarely observed fluorescent activities in the vms, HSNs, and VCs in the absence of the EF stimulus, which shows the sensitivity of these cells to EF (Supplementary Video S6, S8, and S10). That was also expected due to the use of M9 buffer, a hypertonic solution, which inhibits the worm from egg-deposition^[22]^, proving that the proposed electric egg extraction technique is solely stimulated through the application of DC EF. According to these results, we anticipate that our system will be of interest to further investigate the electrosensation in *C. elegans*.

For population investigations, we overlapped the cycles associated with anodal exposures with and without egg-deposition, as well as cathodal exposures (all without egg deposition) for the vms, HSNs, and VC neurons (Figure 9A-C). Fluorescent images of representative worms are also provided prior, during, and after an EF pulse stimulation for all cases. The overlapped cycles showed robust Ca^2+^ transients in vms, HSNs, and VC neurons when worms were facing the anode regardless of the egg-laying event. In contrast and when the worms were facing the cathode, a slight increase in the Ca^2+^ transients during the EF with a small peak right after the EF pulse was observed in vms, while no response was given by HSNs and VC neurons in this condition. These findings show that the cell activities are associated with the EF direction and egg-laying happens only when the HSNs, VC neurons and vms are strongly stimulated in the anodal mode. Our findings also suggest that the egg-laying circuit is involved in *C. elegans* electrosensation, and perhaps electrotaxis, which has not been reported previously.

**Figure 9:**
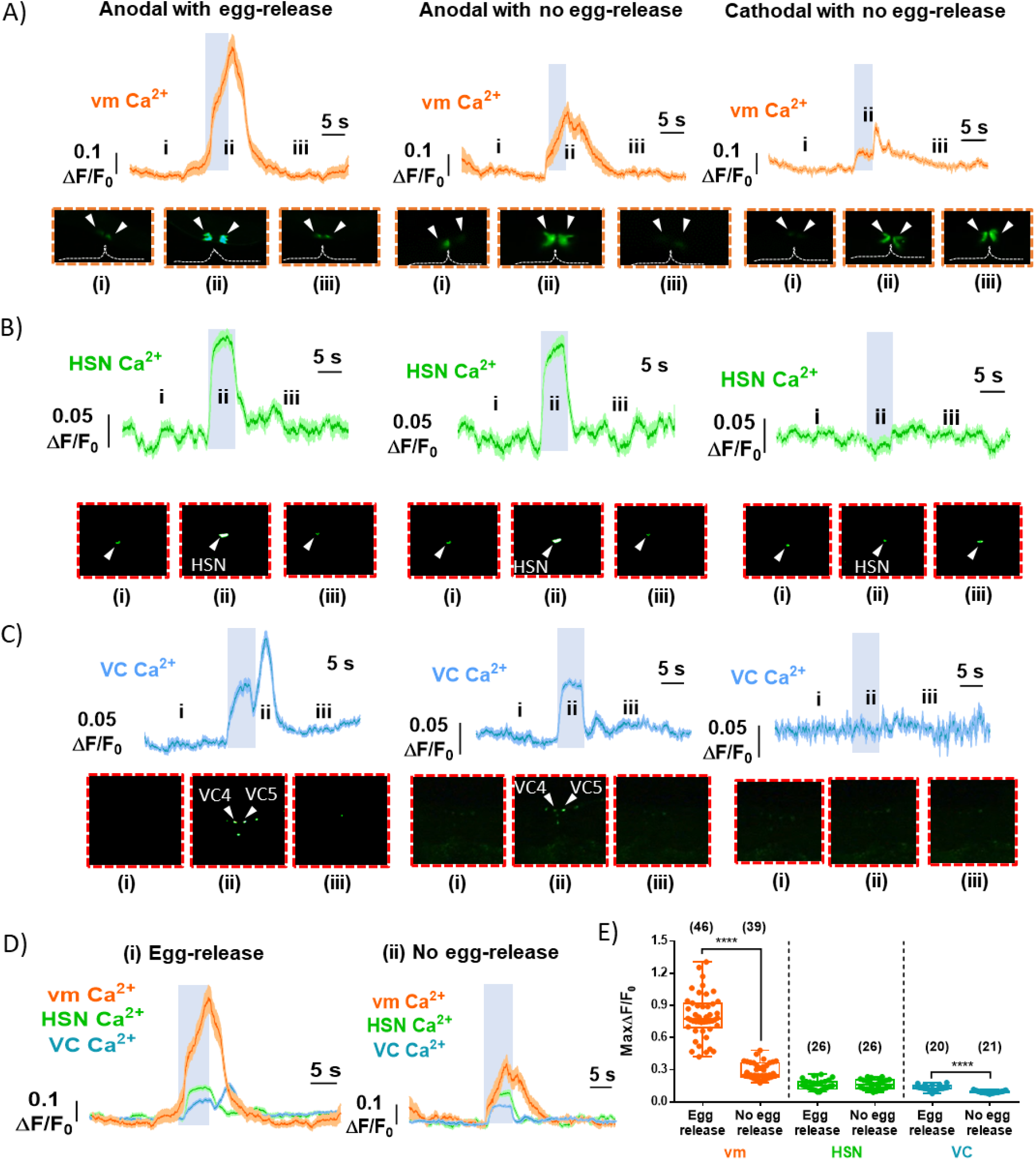
Ca^2+^ transients and individual fluorescence imaging frames recorded during anodal stimulation (left with egg-release and middle with no egg-release) and cathodal stimulation (right with no egg-release) for (A) vms, (B) HSNs, and (C) VC neurons. (D) Overlapped Ca^2+^ transients of the vms, HSN, and VC neurons during (i) egg-release and (ii) no egg-release states. (E) Peak normalized fluorescent activities of vms, HSNs and VC neurons during egg release and no egg release states under anodal stimulation. In all conditions, 6 V/cm EF pulses with 5s on and 25s off were used.

Figure 9D shows the overlapped transient activities of vms, HSNs and VC neurons during anodal exposure, separately illustrated for egg-release or no egg-release states. We noticed that egg-laying usually happened after HSN activation during the EF pulse, which was not different between the egg-release and no egg-release states. This suggests that HSNs activation is necessary for the worm to enter the egg-laying active state which matches with the results obtained by Collins et al.^[1]^. VC neurons and vms were also activated but with different patterns based on egg release. Mechanical deformation of the vulva happening due to egg-deposition was correlated with a significant increase in the Ca^2+^ transients of both the vms during the pulse and the VC neurons after the pulse.

In order to quantitatively determine the cell excitation levels for each strain, we plotted the peak intensities at anodal stimulation during the egg-release and no egg-release states (Figure 9E). vms twitches were observed strongly during the egg-laying events with a wide opening of the vulva to allow for egg-deposition (Figure 9A), whereas the no-egg laying events were associated with significantly weaker twitches (p<0.0001) (Figure 9A and E). The peak anodal responses of HSNs were statistically similar in amplitude during egg release or no egg release events (Figure 9E). VC peak transients were similar in amplitude to the HSNs and were mostly related to the VC4 and VC5 that are in proximity to the vulva (Figure 8A). We noticed an increase in the VCs Ca^2+^ transients during the egg-release compared to no egg release state which were statistically dissimilar (P<0.0001, Figure 9E). We attributed the increase in the activities of vms and VC neurons, during the egg-laying states, to the actual deformation caused by the egg while passing through the vulva, which has also been suggested by Collins et al.^[1]^ This study has provided exciting insights about the egg-laying pathway due to exposure to EF, but the reason behind cell excitability towards the anode, not the cathode, needs further investigation.

### 3.7. Electric egg-laying behaviour of HSN mutant C. elegans

HSNs, located in the vulva area, play a vital role in the egg-laying process^[21]^. Vulva muscles are directly connected to the HSNs through neuromuscular junctions. A substantial reduction in the egg-laying rate has been associated with the ablation or abnormal development of the HSNs ^[1,21,22]^. HSNs mutations have contributed to the understanding of the rhythmic behaviour of egg-laying and it has been shown that exogenous serotonin can directly stimulate the vms to lay eggs in the absence of HSNs^[1]^. In the previous section, we showed that HSNs are activated during the anodal EF excitation period to stimulate the vms for egg release. Here, we aimed at investigating whether the loss of HSNs will reduce the electric egg-laying rate to show the application of our technique in mutant screening.

Age synchronized MT1082 worms (*egl-1*) were tested at the same age as the wildtype N2 worms (D1 past young adult stage). Normally, the MT1082 strain is strongly defective in egg-laying with a bag of ~50 eggs accumulated in their uterus compared to ~15 eggs in the wildtype (Figure 10A) and a resultant larger size (Figure 10B) (Supplementary Video S11). The egg-laying rate and swimming kinematics of the MT1082 in response to 6 V/cm EF pulses (5s on and 25s off) were investigated in comparison to the N2 strain with the results shown in Figure 10C.

**Figure 10:**
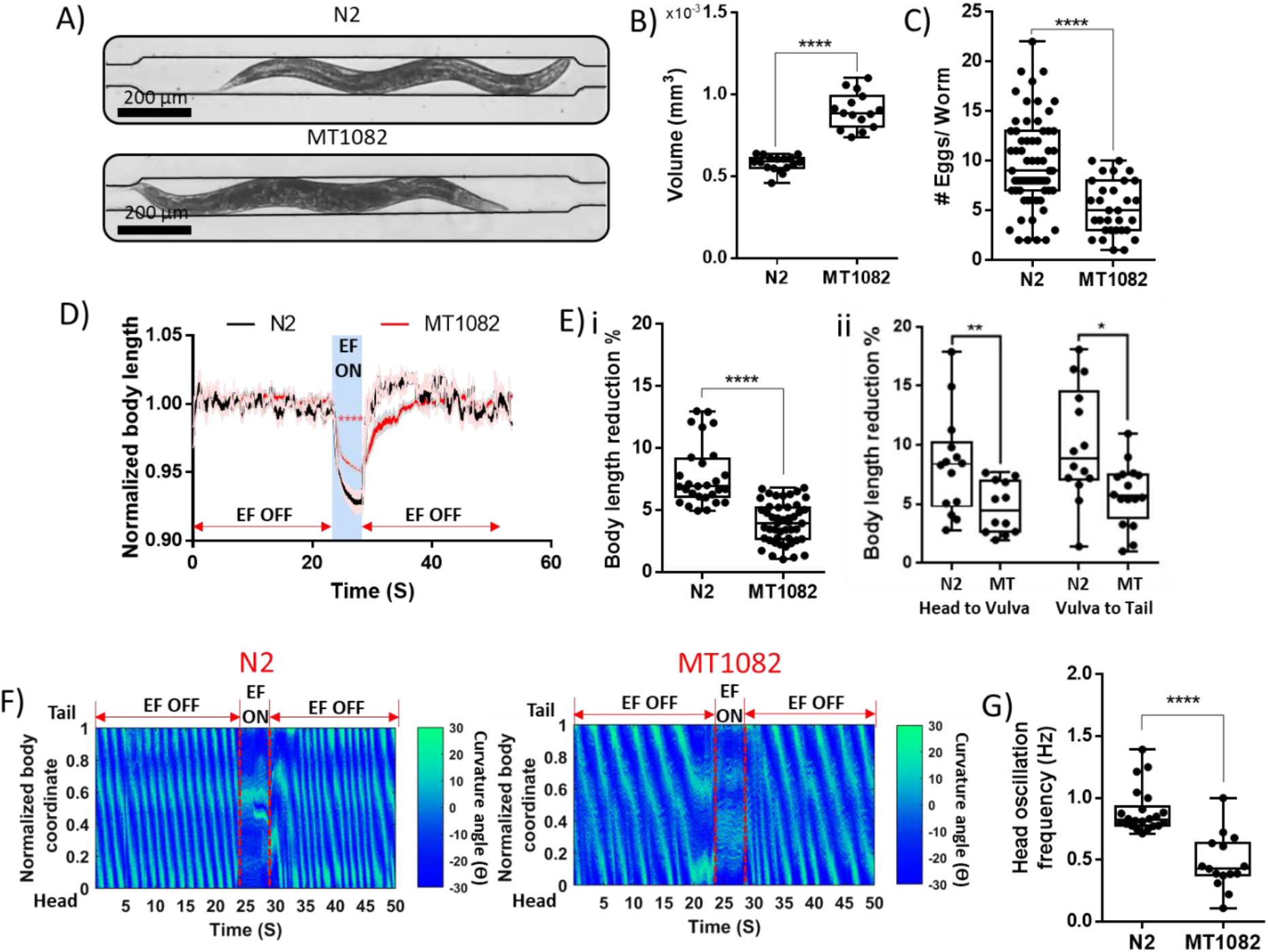
The effect of EF on the worms egg-laying behaviour for wildtype N2 and MT1082, supported by quantitative analysis of the worm kinematics. EF of 6 V/cm was applied for up to 10 minutes in pulses with 5s on and 25s off cycles. (A) Bright-field images of both strains in the microfluidic device, showing that MT1082 contains more eggs (bloating behaviour) at the age of D1. (B) Approximated volume comparison between the two strains. (C) Number of eggs per worm for N = 65 worms for N2 and N= 31 worms for MT1082. (D) Instantaneous normalized body length of both strains (N=18). (E) (i) Maximum total body length reduction and (ii) head-to-vulva and vulva-to-tail length reduction of both strains. (F) Time-lapse normalized body curvature angle of N2 and MT1082. (G) Worms head oscillation frequency during the off-EF periods.

Our results in Figure 10C showed that, out of a total of 100 worms tested, N=31 of MT1082 responded to the EF and expelled a significantly less number of eggs per worm compared to the N2 (N= 65 responders). It is worth emphasizing that the number of non-responders to EF in the MT1082 was significant which reflects the fact that MT1082 is strongly defective in egg-laying even in the presence of EF. As anticipated from our previous results, the decrease in egg-laying was accompanied by a decrease in the body length shortening of MT1082 compared to the N2, as shown in Figure 10D. Moreover, the reduction in length was more towards the tail than the head, like the wildtype behaviour (Figure 10E). Figure 10F shows the curvature contours of both strains for a one-pulse stimulation cycle. The swimming frequency of MT1082 was significantly lower than the N2 during the off-EF period, quantified in Figure 10G, whereas both strains completely stopped movement during the on-EF period. Altogether, our microfluidic chip has aided, for the first time, in studying the role of HSNs in electrosensation and egg laying, and we showed that loss of HSNs does not fully stop the electrical egg deposition, but it decreases the egg-laying rate significantly.

## 4. CONCLUSION

Like many model organisms, *C. elegans* respond to the electric field (EF) in a robust, concise, and persistent manner, by crawling towards the negative electrode of a polarized system, a behaviour termed electrotaxis. Previous studies have solely focused on the electrotactic-induced *C. elegans’* movement and its use for chemical and genetic screening within microfluidic devices. In this paper, we further investigated *C. elegans* electrosensation and introduced, for the first time, a novel EF-evoked behaviour, termed electric egg-laying in a simple to use microfluidic device that enabled trapping and exposure of individual worms to controlled EF conditions. We investigated the effect of worm aging and EF strength, direction, and pulse duration on the egg-laying behaviour, and showed that on-demand egg-deposition could be electrically stimulated in our microfluidic device.

Briefly, we characterized the electric egg-laying behaviour in a worm-fitting microchannel and determined the limiting worm age and EF strength, direction, and pulse duration within which maximum egg-deposition is observed. Egg-count was quantified along with several other phenotypes, including body length, head movement, and transient neuronal activities as measures of electrosensation at a single animal resolution. Interestingly and similar to stimulation of electrotaxis behaviour, electric egg-laying was maximized for the anode-facing worms with a significant increase in the egg-count and reduction in the body length and head motion. The EF stimulation was found to be associated with shortening in the body length and a decrease in the head movement frequency. Moreover, the worms appeared significantly more sensitive at the EF strength of 6V/cm, while all worms exhibited significantly lower response at low EF strengths. The egg-count was found to be independent of pulse duration, whereas the egg-laying response rate and duration were indirectly proportional to pulse duration. Due to the reduction in response duration at longer pulses, our technique has the potential to provide the pathway for developing a high-throughput egg synchronization method without involvement of chemicals (e.g. bleach) or harming the organism.

Our findings also implied that electric egg-laying decreases with worm aging, following the natural off-chip behaviour. Previous studies have shown that egg-laying can be used as a quantitative and user-independent indicator for drug screening and physiological aging. However, aging and life-span studies utilize Fluorodeoxyuridine (FUdR), an inhibitor of DNA synthesis, in a *C. elegans* population to block progeny production and assist the maintenance of synchronous nematode populations for longer times^[34]^. FUdR has shown complex effects on aging in *C. elegans*, which opposes its use in aging studies. This gap in experimental procedures could be breached through the use of EF induced egg-release to deplete the uterus allowing for an extended period without progeny production. Moreover, egg-laying has been reported as a quick technique for antidiabetic drug screening [34]. Therefore, our technique can be an asset to shortening the egg-laying rate assay time and provide a fast quantification technique for drug screening. We anticipate that an appealing follow-up would be to validate the technique using established off-chip protocols for disease-specific compounds.

This is the first time a connection between the egg-laying circuit and electrosensation is reported, unraveling a pathway for further investigation and mapping of the neural basis governing this phenomenon in *C. elegans*. Various strains, expressing intracellular calcium ion dynamics among the egg-laying neural circuit, including HSNs, VC neurons, and vms were imaged during the EF stimulation. The results exhibited EF directional-dependency of the HSNs, VCs, and vms, which might elucidate the innate cathode-driven orientation of *C. elegans* as a safety mechanism to lower the excitation of its cells. We further tested a mutated strain lacking the HSNs and demonstrated that EF still induces egg-laying but with a defective phenotype, which shows the applicability of our technique for mutant screening. We envision that with small modifications to the existing design, several nematodes can be tested simultaneously under identical conditions, accommodating the needs to develop higher throughput electric egg-laying screening devices.

Altogether, our paper discusses novel electrically induced physiological and behavioural phenotypes discovered in *C. elegans* aiming to drive new hypotheses on the biological mechanisms governing electrosensation. Additionally, we established that EF stimulation of *C. elegans* within our microfluidic device has no adverse effects on the physiology of the nematodes.

## ACKNOWLEDGEMENTS

This work was supported by Natural Sciences and Engineering Research Council (NSERC) of Canada and the Early Researcher Award to PR and the Ontario Trillium Scholarship to KY.

## References

[1] K. M. Collins, A. Bode, R. W. Fernandez, J. E. Tanis, J. C. Brewer, M. S. Creamer, M. R. Koelle, eLife 2016, 5, e21126.

[2] A. C. Hart, 2006.

[3] B. P. Gupta, P. Rezai, Micromachines 2016, 7, 123.

[4] H. E. Kinser, Z. Pincus, Molecular and Cellular Neuroscience 2017, 80, 192.

[5] A. J. Harrington, S. Hamamichi, G. A. Caldwell, K. A. Caldwell, Developmental Dynamics 2010, 239, 1282.

[6] C. D. Link, Experimental Gerontology 2006, 41, 1007.

[7] M. Artal-Sanz, L. de Jong, N. Tavernarakis, Biotechnology Journal 2006, 1, 1405.

[8] K. Youssef, A. Tandon, P. Rezai, Integrative biology: quantitative biosciences from nano to macro 2019, 11, 186.

[9] K. Youssef, P. Bayat, A. R. Peimani, S. Dibaji, P. Rezai, in Environmental, Chemical and Medical Sensors, Springer, 2018, pp. 199–225.

[10] M. M. Shanmugam, Biology, Engineering and Medicine 2017, 2, 1.

[11] N. C. Sukul, N. A. Croll, Journal of nematology 1978, 10, 314.

[12] C. V. Gabel, H. Gabel, D. Pavlichin, A. Kao, D. A. Clark, A. D. T. Samuel, Journal of Neuroscience 2007, 27, 7586.

[13] S. D. Chrisman, C. B. Waite, A. G. Scoville, L. Carnell, PLoS ONE 2016, 11, DOI 10.1371/journal.pone.0151320.

[14] H. S. Chuang, W. J. Kuo, C. L. Lee, I. H. Chu, C. S. Chen, Scientific Reports 2016, 6, 1.

[15] P. Rezai, A. Siddiqui, P. R. Selvaganapathy, B. P. Gupta, Lab on a Chip 2010, 10, 220.

[16] P. Rezai, A. Siddiqui, P. R. Selvaganapathy, B. P. Gupta, Applied Physics Letters 2010, 96, 153702.

[17] S. Salam, A. Ansari, S. Amon, P. Rezai, P. R. Selvaganapathy, R. K. Mishra, B. P. Gupta, in Worm, Taylor & Francis, 2013, p. e24558.

[18] P. Rezai, S. Salam, B. P. Gupta, P. R. Selvaganapathy, 15th International Conference on Miniaturized Systems for Chemistry and Life Sciences 2011, MicroTAS 2011 2011, 2, 723.

[19] X. Wang, R. Hu, A. Ge, L. Hu, S. Wang, X. Feng, W. Du, B. F. Liu, Lab on a Chip 2015, 15, 2513.

[20] B. Han, D. Kim, U. H. Ko, J. H. Shin, Lab on a chip 2012, 12, 4128.

[21] W. R. Schafer, Annual Review of Genetics 2006, 40, 487.

[22] W. R. Schafer, in WormBook: The Online Review of C. Elegans Biology [Internet], WormBook, 2005.

[23] E. R. Sawin, 1996.

[24] L. A. Fenk, M. de Bono, Proceedings of the National Academy of Sciences 2015, 112, E3525.

[25] T. Stiernagle, C. elegans 1999, 2, 51.

[26] M. Porta-de-la-Riva, L. Fontrodona, A. Villanueva, J. Cerón, Journal of Visualized Experiments 2012, e4019.

[27] Y. Xia, G. M. Whitesides, Annual Review of Materials Science 1998, 28, 153.

[28] J. Schindelin, I. Arganda-Carreras, E. Frise, V. Kaynig, M. Longair, T. Pietzsch, S. Preibisch, C. Rueden, S. Saalfeld, B. Schmid, J. Y. Tinevez, D. J. White, V. Hartenstein, K. Eliceiri, P. Tomancak, A. Cardona, Nature Methods 2012, 9, 676.

[29] F. Faul, E. Erdfelder, A. G. Lang, A. Buchner, Behavior Research Methods 2007, 39, 175.

[30] X. Manière, F. Lebois, I. Matic, B. Ladoux, J.-M. Di Meglio, P. Hersen, PloS one 2011, 6.

[31] H. Hwang, D. E. Barnes, Y. Matsunaga, G. M. Benian, S. Ono, H. Lu, Scientific reports 2016, 6, 19900.

[32] X. Chen, J. W. Barclay, R. D. Burgoyne, A. Morgan, Chemistry Central Journal 2015, 9, 65.

[33] C. L. Pickett, K. Kornfeld, Aging cell 2013, 12, 544.

[34] H. Wang, Y. Zhao, Z. Zhang, Biochemical and biophysical research communications 2019, 509, 694.

